# A Method For Estimating The Cholesterol Affinity Of Integral Membrane Proteins From Experimental Data

**DOI:** 10.1101/2023.10.03.560595

**Authors:** Theodore L. Steck, S. M. Ali Tabei, Yvonne Lange

## Abstract

The cholesterol affinities of many integral plasma membrane proteins have been estimated by molecular computation. However, these values lack experimental confirmation. We therefore developed a simple mathematical model to extract sterol affinity constants and stoichiometries from published isotherms for the dependence of the activity of such proteins on membrane cholesterol concentration. The binding curves for these proteins are sigmoidal with strongly-lagged thresholds attributable to competition for the cholesterol by bilayer phospholipids. The model provided isotherms that matched the experimental data using published values for the sterol association constants and stoichiometries of the phospholipids. Three oligomeric transporters were found to bind cholesterol without cooperativity with dimensionless association constants of 35 for Kir3.4* and 100 for both Kir2 and a GAT transporter. (The corresponding ρG° values were -8.8, -11.4 and -11.4 KJ/mol, respectively.) These association constants are significantly lower than those for the phospholipids which range from ∼100 to 6,000. The BK channel, the nicotinic acetylcholine receptor and the M192I mutant of Kir3.4* appear to bind multiple cholesterol molecules cooperatively (n = 2 or 4) with subunit affinities of 563, 950 and 700, respectively. The model predicts that the three less avid transporters are approximately half-saturated in their native plasma membranes; hence, sensitive to variations in cholesterol in vivo. The more avid proteins would be nearly saturated in vivo. The method can be applied to any integral protein or other ligand in any bilayer for which there are reasonable estimates of the sterol affinities and stoichiometries of the phospholipids.

## INTRODUCTION

Cholesterol is bound to many plasma membrane proteins, affecting their disposition and activity in diverse ways. These associations have been characterized by various techniques, including X-ray, cryo-EM, NMR, molecular computation and functional analysis.^1–10^ Two cardinal features of the interaction of the proteins and sterols are the strength of their association and their binding stoichiometry. We have found only one experimental determination in the literature of the affinity of cholesterol for an integral membrane protein in situ.^11^ However, that study is not secure.^12^ In the absence of experimental values, sterol binding constants for numerous proteins have been obtained by molecular dynamics, docking and other computational methods.^4, 13–21^

Here, we describe an indirect method to characterize the binding of cholesterol to integral proteins and other membrane-embedded ligands. We developed a simple mathematical model that takes advantage of the formation of complexes between cholesterol and membrane phospholipids which, perforce, compete with proteins for the sterol.^22–24^ The sterol dependence of the activity of the proteins is consequently constrained by the sterol affinity and stoichiometry of the membrane phospholipids. Using literature values for the latter, one can obtain estimates of the sterol affinity and stoichiometry of these proteins by matching computer-simulated binding isotherms to experimental cholesterol dependence curves for protein activity. In other words, competition by phospholipids with known cholesterol affinity provide a quantitative gauge of the binding of the sterol to integral membrane proteins.

At one extreme, very avid proteins outcompete the phospholipids for the cholesterol and so bind it with nearly hyperbolic isotherms. At the other extreme, very weak proteins bind cholesterol poorly until the phospholipids become saturated with sterol at their stoichiometric equivalence point and the uncomplexed excess sterol becomes increasingly available to the protein.^25^ Between these extremes, proteins have sigmoidal sterol dependence curves with shapes and thresholds determined both by the ligands and the phospholipids (see Results). We assume that two forms of cholesterol are available to the protein: (a) that extracted from membrane phospholipid complexes by strong competition; and (b) an uncomplexed fraction readily available to the protein beyond the stoichiometric equivalence point of the phospholipid complexes.^25^

## METHODS

### The model

This model simulates the association of sterol molecules with ligands such as proteins within a phospholipid bilayer so as to estimate the affinity and stoichiometry of the binding reaction of the ligand. It resembles an earlier model that treats the binding of water-soluble ligands such as cytolysins to membrane sterols.^25^ In contrast, the present version places the ligand in the same membrane compartment as the sterol and the phospholipid. It utilizes the simplest formalism consistent with the experimental data: competitive binding of sterols to phospholipids constrains their binding to the proteins. Applying values for the former allows values for the latter to be obtained through computed simulations.

We assume that a ligand (L) associates with cholesterol (C) in competition with membrane phospholipids (P). The essential parameters are therefore the sterol association constants and stoichiometries for these reactants. The compartment size is α = (C_T_ + P_T_ + L_T_), where the subscript T denotes total. The chemical activities of the reactants are ideal and expressed as their concentrations in mole fractions. These are C_f_/α, P_f_/α and L_f_/α, where f denotes their free or unbound form. CP_r_ denotes lipid complexes with a stoichiometry of one cholesterol to r phospholipid molecules. C_n_L denotes cholesterol-ligand (e.g., protein) complexes with a stoichiometry of n cholesterol molecules per ligand.

The association equilibrium for the sterol and phospholipid is:

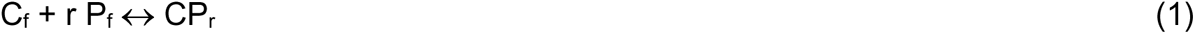

The dimensionless association constant for complexes is then

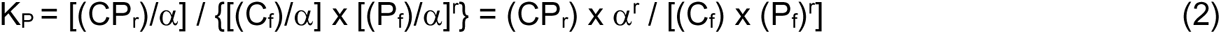

Many integral membrane proteins such as the transporters considered here are oligomeric.^26, 27^ For simplicity, we allow the ligand (L) to be an oligomer composed of identical subunits that bind n sterol molecules in a fully cooperative all-or-none fashion; a concerted rather than sequential reaction.^28^ Hence,

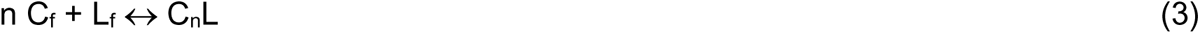

Designating the dimensionless association constant for the formation of an n-mer of L as K_Ln_, we have

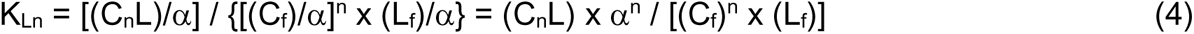

### Application of the model

Eqs.1-4 were used to generate isotherms that simulate the binding of an integral membrane protein to cholesterol in competition with the phospholipids. Simulations are numerical solutions computed in MATLAB. The computer code is presented in the S2 section of the Supporting Information. Reactant quantities are in moles. The abundance of the phospholipid (P_T_) was set to unity and the mole fraction of the sterol (C_T_/α) was varied across its physiologic range, from zero to 60 mol%. The abundance of the protein (L_T_) was set at 1 x 10^-3^ for biological membranes and 1 x 10^-5^ for planar bilayers. (The concentration of the protein affected the results by far less than one percent--nor should competition by extraneous proteins matter because of the large molar excess of the sterol and phospholipids.) Cholesterol/protein binding ratios of n = 1 to 5 were examined.

Accurate values for the sterol affinity and stoichiometry of the phospholipid mixtures (i.e., K_P_ and CP_r_ in Eq. 2) were not available. We therefore used plausible estimates based on a published data set.^29^ We assumed that the stoichiometry of the binding of phospholipid to cholesterol, r, was one or two.^24, 25, 29^ The program allows membranes to contain two types of phospholipids with different cholesterol affinities and stoichiometries in different proportions. The membrane sterol concentration was expressed as mol% cholesterol; *i.e.*, 100 x moles cholesterol / (moles cholesterol + moles phospholipid).

Assuming plausible values for the cholesterol affinity and stoichiometry of the phospholipids, we tested a range of values for each protein by matching computed isotherms to the experimental data by eye. The experimental data were kept in their published form, scaled on the left-hand ordinates of the figures. No data points were omitted. Simulated isotherms were scaled from zero to one on the right-hand ordinates. In addition to the cholesterol binding affinity and stoichiometry of a protein, K_Ln_ and n, the simulations yielded cholesterol-dependence curves for the fractional saturation of the protein (C_n_L/L_T_), the fraction of each phospholipid associated with the sterol (r x CP_r_/P_T_) and the fraction of uncomplexed sterol (C_f_/C_T_). Free energies of sterol binding were calculated as ΔG° = -RT ln K_L1_ at 298 °K.

A satisfactory match of simulation to data was found in each case. However, estimates of K_Ln_ depended on K_P_. Consequently, the experimental data could be matched by various pairs of values for K_P_ and K_L_. Thus, the delivered association constants were not uniquely determined and were therefore not definitive.

It is shown in Section S3.a of the Supporting Information that, at half-saturation of the protein,

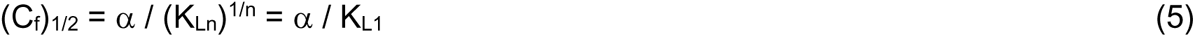

Eq. (5) implies that the half-saturation of a protein is independent of the assumed sterol binding stoichiometry, n, and, furthermore, that the cholesterol-dependent isotherms of a protein will all intersect at (C_f_)_1/2_ (not shown).

In addition, note that the calculated isotherms in each of several figures intersect close to the same *total* cholesterol concentration (C_T_) for all values of n. As shown in Section S3.b of the Supporting Information, this intersection point is given by:

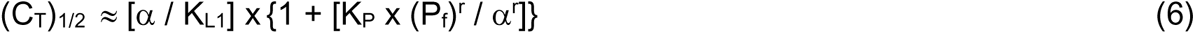

Thus, the membrane concentration of cholesterol at the half-saturation point of a protein with affinity K_Ln_ is essentially independent of n.

## RESULTS

Many integral plasma membrane proteins associate with cholesterol (see Introduction). However, sterol-dependent activity curves suitable for our analysis are few.^30^ Six such isotherms were tested here.

### GAT

These oligomeric transporters facilitate the recovery of secreted γ-aminobutyric acid (GABA) across presynaptic plasma membranes, symported by gradients of NaCl.^31^ The data in Figure 1 show the cholesterol dependence of the transport activity of a purified rat brain GAT reconstituted in liposomes.^32^ The overall sterol affinity of the phospholipids employed was not known, so we tested some relevant K_P_ values.^29^ A very good match to the data was obtained by assuming an affinity of the phospholipid for the sterol of K_P_ = 200, a protein affinity of K_L1_ = 100 and a protein stoichiometry of n = 1 (curve 3 in Figure 1). Because the protein competes moderately well with the phospholipids, the isotherm has a gentle sigmoidicity and ascends well before the appearance of free cholesterol at its presumed stoichiometric equivalence point with the phospholipids, 43 mol% (curve 6). This behavior shows that the protein extracts cholesterol from the phospholipid complexes.

**Figure 1.**
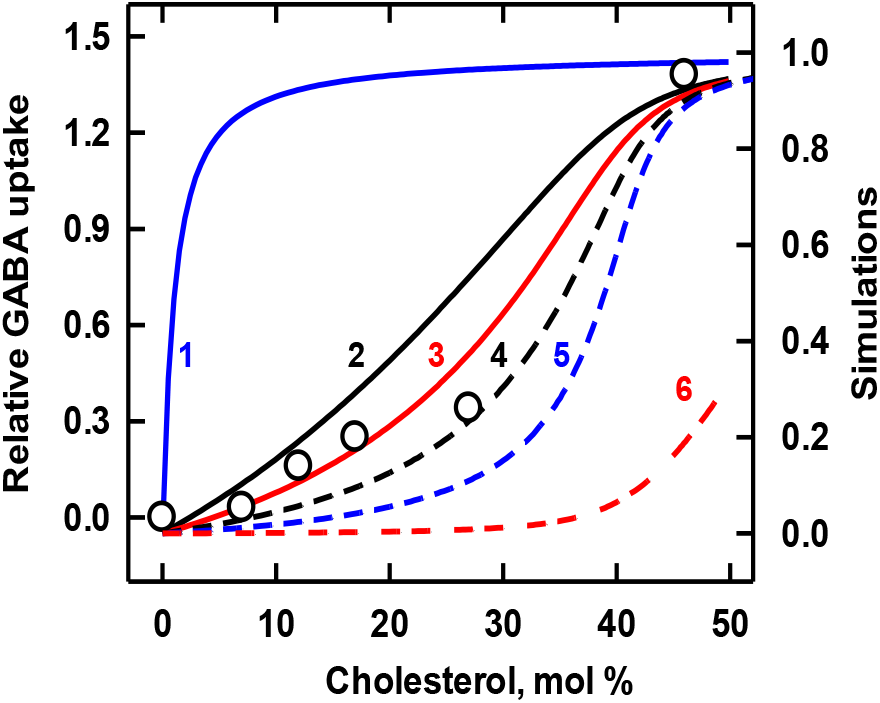
Simulation of the binding of cholesterol to a GAT transporter. Experimental data (symbols; taken from Figure 4 in ref. 32) depict the uptake of γ-aminobutyric acid (GABA) by liposomes containing the transporter, asolectin, brain phospholipids and varied cholesterol.^32^ Curves 1-5 assigned K_L1_ = 100 and n = 1 to the protein. Lacking experimental values for the phospholipid mixture, we assumed that the bilayer contained equal proportions of two phospholipid species with stoichiometries of C:P = 1:1 and 1:2, with K_P_ association constants shown in Table 1.^29^ The stoichiometric equivalence point of such bilayers is 43 mol% cholesterol. Curve 6 shows free cholesterol in moles/mol phospholipid in the absence of protein.

**Table 1.**
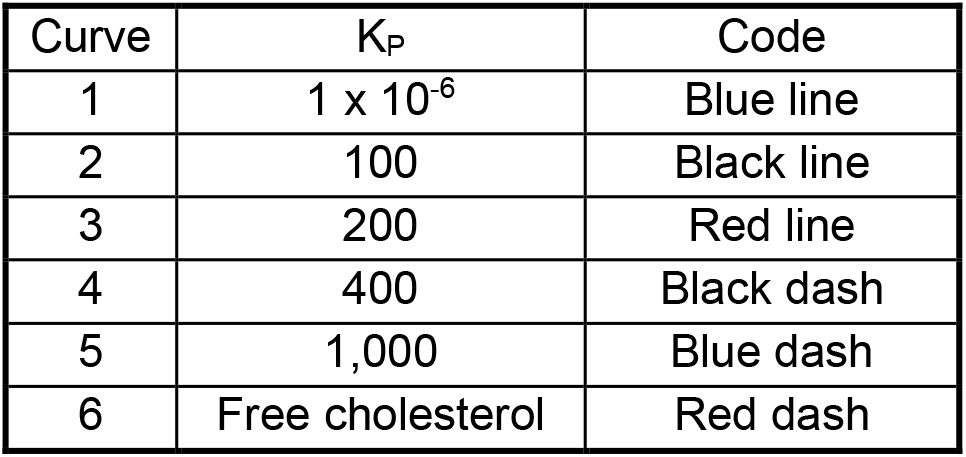
Values used for GAT in Figure 1.

| Curve | $K_P$ | Code |
| --- | --- | --- |
| 1 | $1 \times 10^{-6}$ | Blue line |
| 2 | 100 | Black line |
| 3 | 200 | Red line |
| 4 | 400 | Black dash |
| 5 | 1,000 | Blue dash |
| 6 | Free cholesterol | Red dash |

The other curves in Figure 1 explore the model. Curve 1 shows that, in the presence of phospholipids with low sterol affinity, avid proteins will bind cholesterol strongly with a nearly hyperbolic isotherm. Curves 2 and 4 show the effect of doubling and halving the sterol affinity of the best-matched phospholipids (curve 3). The spread among curves 2, 3 and 4 gives an indication of the sensitivity to phospholipid affinity of the sterol-protein interaction. Curve 5 illustrates how the cholesterol binding curve of a protein is shifted to the right when the bilayer phospholipids are very avid.

We also matched the experimental data to simulations that assumed that the protein has two or four identical sterol binding sites; that is, n = 2 and 4. The affinity of the monomer was again taken to be K_L1_ = 100. The curves had lagged thresholds and very acute ascents and so did not match the experimental data well (not shown). We infer that the binding of cholesterol to the GAT is not cooperative. Nevertheless, multiple cholesterol molecules might associate with the oligomer at non-interacting sites.

### KIR3.4*

This recombinant homotetramer was derived from a subunit of the G protein-activated potassium channel, GIRK.^8^ In the study shown in Figure 2, the protein was reconstituted into planar bilayers and single channel activities were recorded as a function of membrane cholesterol concentration.^33^ Curve 2 provides a good match to the data using K_L1_ = 35 and n = 1. Doubling and halving the sterol affinity of the simulated protein (curves 1 and 3, respectively) shifted the binding isotherm moderately. Curve 4 assumed n = 2; it paralleled the data at low cholesterol concentrations but rose acutely and overshot the plateau. Hence, we preferred the match for n = 1. For illustrative purposes, curve 5 shows a protein with a weak affinity that nevertheless extracts cholesterol from the phospholipids. Curve 6 shows the concentration of uncomplexed cholesterol in the absence of protein; it rises acutely as the total cholesterol exceeds the complexation capacity of the phospholipids at their stoichiometric equivalence point, 43 mol%.

**Figure 2.**
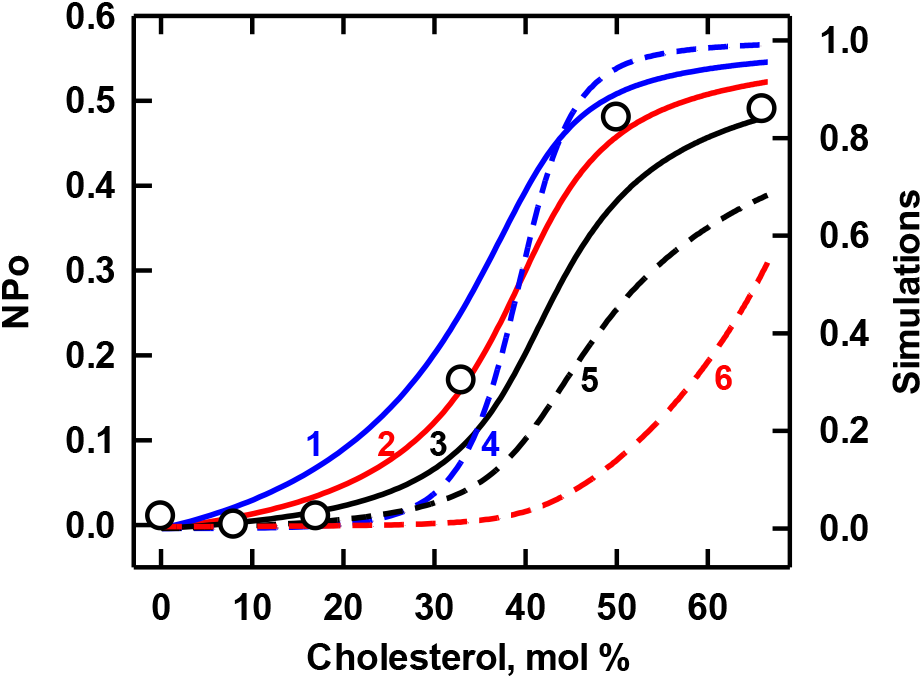
Simulation of the binding of cholesterol to Kir3.4*. Experimental data (symbols; taken from Figure 1H in ref. 33) represent the open probability (NPo) of single channels in a planar bilayer composed of 1:1 mixtures of 1-palmitoyl-2-oleoyl-phosphatidylserine (POPS) and brain phosphatidylethanolamine (POPE).^33^ The former phospholipid was assigned a sterol stoichiometry of C:P = 1:2 and an affinity of K_P_ = 210; the latter species was assigned a C:P = 1:1 and a K_P_ = 130 (the value published for 1,2-dimyristoyl-phosphatidylethanolamine, DMPE).^29^ The stoichiometric equivalence point of such bilayers is 43 mol% cholesterol. The values for the protein in Table 2 were used in the simulation of the isotherms. Curve 6 shows the concentration of free cholesterol in moles/mol phospholipid in the absence of protein. As predicted by the model, curves 2 and 4 intersect at half-saturation (see Methods).

**Table 2.** Values used for Kir3.4* in Figure 2.

| Curve | n | $K_{Ln}$ | Code |
| --- | --- | --- | --- |
| 1 | 1 | 70 | Blue line |
| 2 | 1 | 35 | Red line |
| 3 | 1 | 17.5 | Black line |
| 4 | 2 | 1,225 | Blue dash |
| 5 | 1 | 5 | Black dash |
| 6 | Free cholesterol |  | Red dash |

### Kir2

As seen in Figure 3, the activity of this tetrameric inward rectifier potassium channel is inhibited by cholesterol.^34, 35^ The simulation of these data is considered to be slightly better for n = 1 (curve 1) than for n = 4 (curve 2), both by eye and from the value of R^2^ (Table 3). Curves for n = 2 and 3 fell between these two extremes (not shown). It appears that the protein does not extract appreciable cholesterol from the avid phospholipid complexes but depends on the uncomplexed cholesterol emerging at the stoichiometric equivalence point of the plasma membrane. This undermines the estimation of the stoichiometry, n, which depends on the shape of the isotherm.

**Figure 3.**
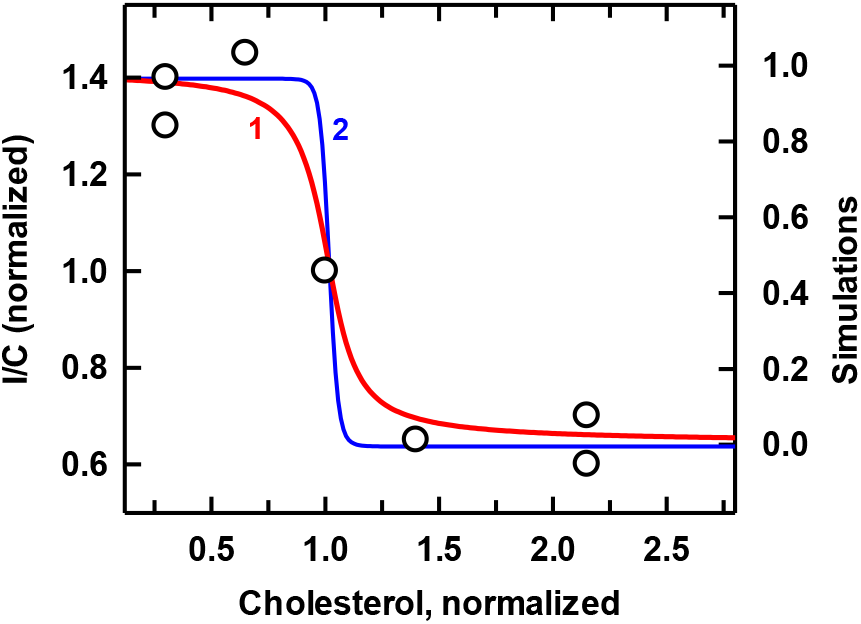
Simulation of the binding of cholesterol to Kir2 channels. Experimental data (symbols; taken from Figure 5 in ref. 34) give the peak current density (normalized I/C) obtained by patch clamping cultured bovine aortic endothelial cells that had been treated with methyl-β-cyclodextrin plus varied cholesterol to adjust the plasma membrane sterol.^34^ The plasma membrane was represented by a mixture of equal proportions of two phospholipids with stoichiometries of C:P = 1:1 and 1:2 and affinities of 5,000 and 2,500, respectively.^24, 25^ The resting plasma membrane cholesterol concentration, here normalized to 1.0, was therefore 43 mol% (mole fraction = 0.43 or C/P = 0.75).^24^ The simulations used the protein values given in Table 3. As predicted by the model, the curves intersected at their half-saturation values (see Methods). [Much of the excess sterol loaded into the cells presumably moved to intracellular organelles.^36^ However, this should not have affected the derived K_Ln_.]

**Table 3.**
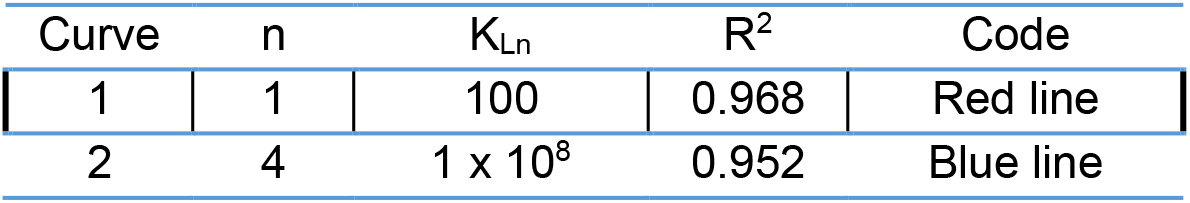
Values used in Kir2 in Figure 3.

### BK

This voltage-gated potassium channel is activated by cytoplasmic calcium ions with high positive cooperativity.^37, 38^ However, as shown in Figure 4, it is inhibited by cholesterol.^38, 39^ The simulation for n = 4 (curve 2; R^2^=0.73) was better matched to the data than that for n = 1 (curve 1; R^2^ = 0.58). Rejecting the stray point at ∼20 weight % further favored the match for n = 4 over that for n = 1 (R^2^ = 0.90 vs. 0.66). The sterol affinity of a putative subunit was K_L1_ = 563 (Table 4). Thus, BK was much more avid for cholesterol than the three transporters analyzed above.

**Figure 4.**
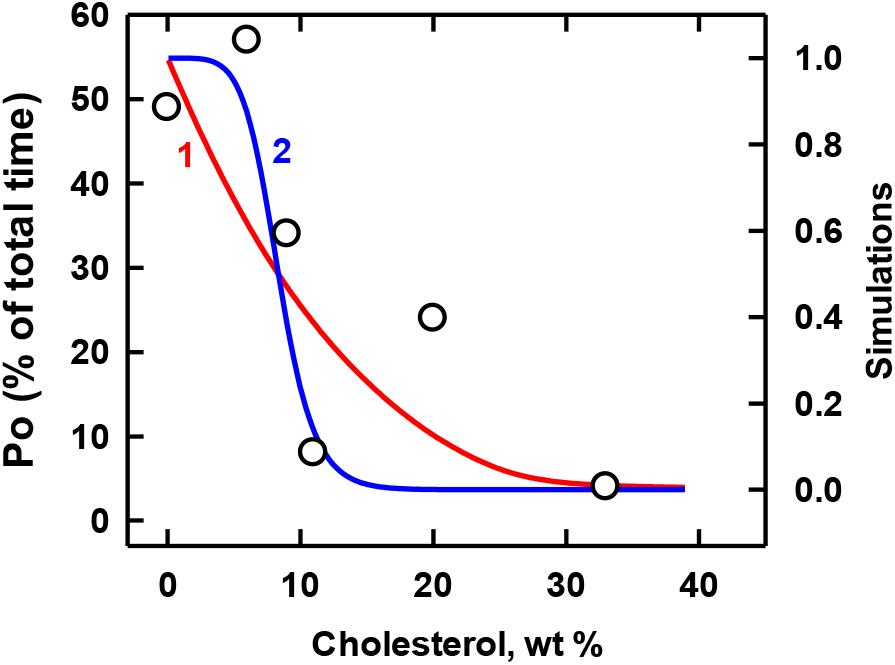
Simulation of the binding of cholesterol to BK channels. (These were presumably tetramers of the alpha subunit.^40^) Experimental data (symbols; taken from Figure 4 in ref. 39) represent the open probability (P_o_) of single channels in a planar bilayer composed of a 1:1 mixture of 1-palmitoyl-2-oleoyl-phosphatidylserine (POPS) plus 1-palmitoyl-2-oleoyl-phosphatidylethanolamine (POPE) in the presence of 0.1 mM CaCl_2_.^39^ We assumed that the two phospholipids had stoichiometries of C:P = 1:2 and 1:1 and affinities of K_P_ = 210 and 130, respectively.^29^ The stoichiometric equivalence point of such bilayers is 43 mol% cholesterol. To convert the experimental cholesterol concentrations from the reported percent of total lipid weight to mol%, we assumed molecular weights of 387 and 740 for C and P, respectively. The midpoint of the curve was 8.3 weight % which corresponds to 12.7 mol% cholesterol. The protein values for BK listed in Table 4 were used in the simulations. As predicted by the model, the curves intersect at their half-saturation values (see Methods).

**Table 4.**
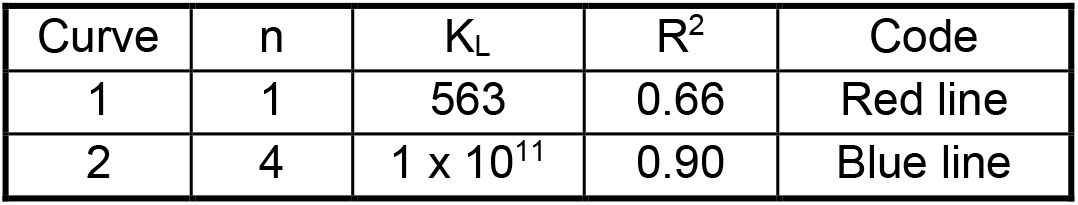
Values used for BK in Figure 4.

| Curve | n | $K_L$ | $R^2$ | Code |
| --- | --- | --- | --- | --- |
| 1 | 1 | 563 | 0.66 | Red line |
| 2 | 4 | $1 \times 10^{11}$ | 0.90 | Blue line |

### M192I variant of Kir3.4*

This mutant recombinant homotetrameric potassium channel was recently characterized.^41^ The cholesterol dependence of its conductance was altered by the mutation in three ways. First, while cholesterol activated the wild type (Figure 2), it inhibited the variant (Figure 5). Second, whereas the open probability of the unmodified channel in Figure 2 appeared to have first-order dependence on cholesterol concentration, the data for the mutant were better matched by assuming n = 4 rather than n = 1. Third, the mutant protein was far more avid for cholesterol than was the wild type (i.e., K_L1_ = 700 versus 35, respectively). Consequently, its sterol threshold was shifted far to the left. Figure 1 shows that the sterol affinity of the phospholipid strongly influences the binding isotherm of a protein. But, given that the Kir3.4* and the M192I variant were analyzed in the same bilayer environment, the striking difference in their isotherms can be ascribed to the alteration in the protein.

**Figure 5.**
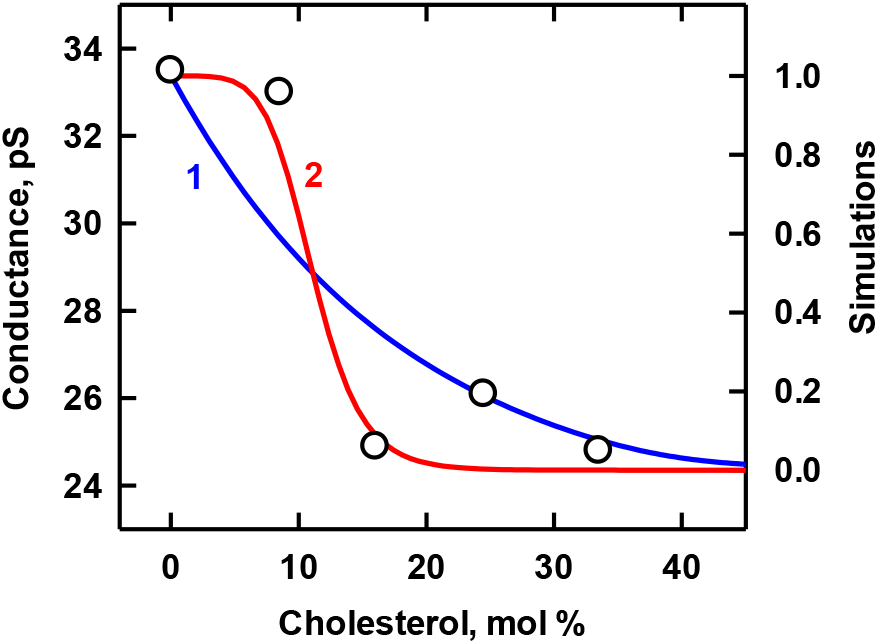
Simulation of the binding of cholesterol to Kir3.4* M182I channels. Experimental data (symbols; taken from Figure 4D in ref. 41) represent the conductance of single channels in a planar bilayer composed of a 1:1 mixture of 1-palmitoyl-2-oleoyl-phosphatidylserine (POPS) and 1-palmitoyl-2-oleoyl-phosphatidylethanolamine (POPE).^41^ We assumed that the two phospholipids had C:P stoichiometries of 1:2 and 1:1 and affinities of K_p_ = 210 and 130, respectively.^29^ The stoichiometric equivalence point of such bilayers is 43 mol% cholesterol. The computed curves used the protein values listed in Table 5. As predicted by the model, the curves intersect at their half-saturation values (see Methods).

**Table 5.** Values used for Kir3.4* M182I in Figure 5.

| Curve | n | $K_L$ | $R^2$ | Code |
| --- | --- | --- | --- | --- |
| 1 | 1 | $7 \times 10^2$ | 0.77 | Blue line |
| 2 | 4 | $2.4 \times 10^{11}$ | 0.93 | Red line |

### Nicotinic acetylcholine receptor (AChR)

This pentameric cation channel has complex allosteric behavior and specific binding sites for cholesterol.^42, 43^ The data in Figure 6 show the sterol-dependence of a conformational switch in the receptor from a desensitized to a resting state.^44^ Simulation with n = 1 (curve 1) gives a poor match to the data, and the threshold for n = 5 (curve 3) is too sharp. On the other hand, assuming modest cooperativity (i.e., n = 2, curve 2) provides an excellent match. Calculations using n = 3 or n = 4 give intermediate simulations while varying the affinity of the phospholipids moves these cooperative curves to lower and higher thresholds as in Figure 1 (not shown). The threshold of the curve was far to the left of that predicted for the phospholipids. Thus, both the cooperativity of the protein and competition with the phospholipids contributed to the position and contour of curve 2, the best-matched simulation.

**Figure 6.**
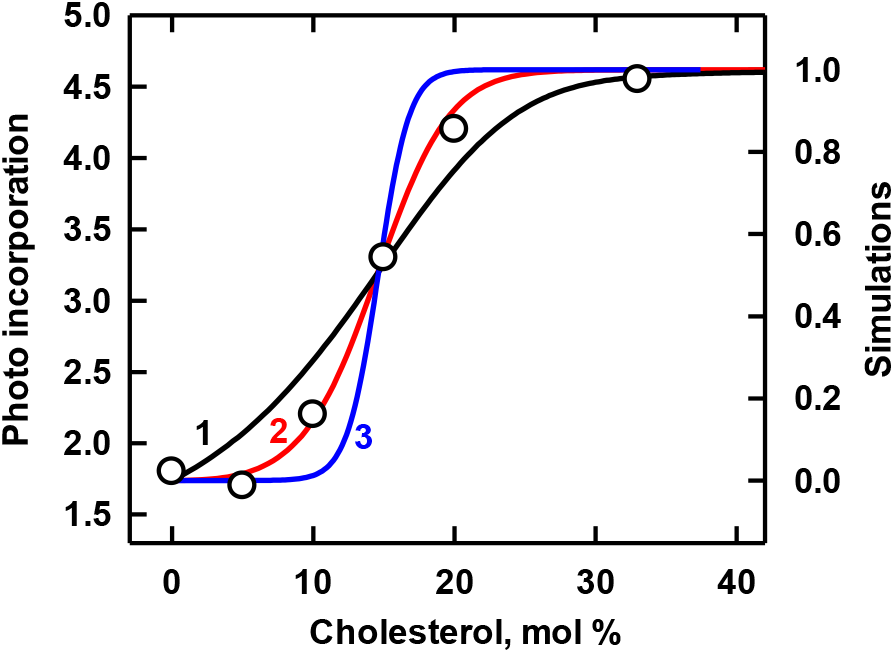
Simulation of the binding of cholesterol to nicotinic acetylcholine receptors. Experimental data (symbols; taken from Figure 1C in ref. 44) represent the labeling of the protein by a conformationally-sensitive radioactive diazirine probe in vesicles composed of 3:1 dioleoylphosphatidylcholine:dioleoylphosphatidic acid (DOPC:DOPA).^44^ We assumed sterol affinities of K_P_ = 930 and 200, respectively, for the two phospholipids and a C:P stoichiometry of 1:2 for both.^29^ The stoichiometric equivalence point of such bilayers is 33 mol% cholesterol. The simulations used the protein values listed in Table 6. As predicted by the model, the curves intersect at their half-saturation values (see Methods).

**Table 6.** Values used in Figure 6.

| Curve | n | $K_L$ | Code |
| --- | --- | --- | --- |
| 1 | 1 | $9.5 \times 10^2$ | Black |
| 2 | 2 | $9 \times 10^5$ | Red |
| 3 | 5 | $7.7 \times 10^{14}$ | Blue |

The sterol affinity of the protein subunit, K_L1_ = 950, is an order of magnitude greater than that of the proteins shown in Figures 1-3. The corresponding free energy change is ρG° = -17.2 KJ/mol. An earlier molecular dynamics simulation suggested an energy of interaction of cholesterol with subunits of the receptor of -25 to -50 KJ/mol.^45^ However, those determinations were performed in vacuo rather than in a bilayer.

### Simulated isotherms for five transporters in cell membranes

The model posits that the value for K_Ln_ does not vary with the phospholipid environment of a protein. This allowed us to apply the K_Ln_ values obtained above to model five of those proteins in a plasma membrane bilayer (Figure 7). (We did not include the M192I mutant of Kir3.4* from Figure 5 because its biological relevance is uncertain.) The midpoints of the simulations for the three low-affinity transporters (curves 3, 4 and 5 in Figure 7) fell near the resting cholesterol concentration assumed for the plasma membrane, ∼43 mol%.^24^ In contrast, the two high-affinity transport proteins with n = 4 (curve 2) and n = 2 (curve 1) had thresholds to the left of the physiological rest point; this predicts that they are saturated with the sterol in the resting plasma membrane.

**Figure 7.**
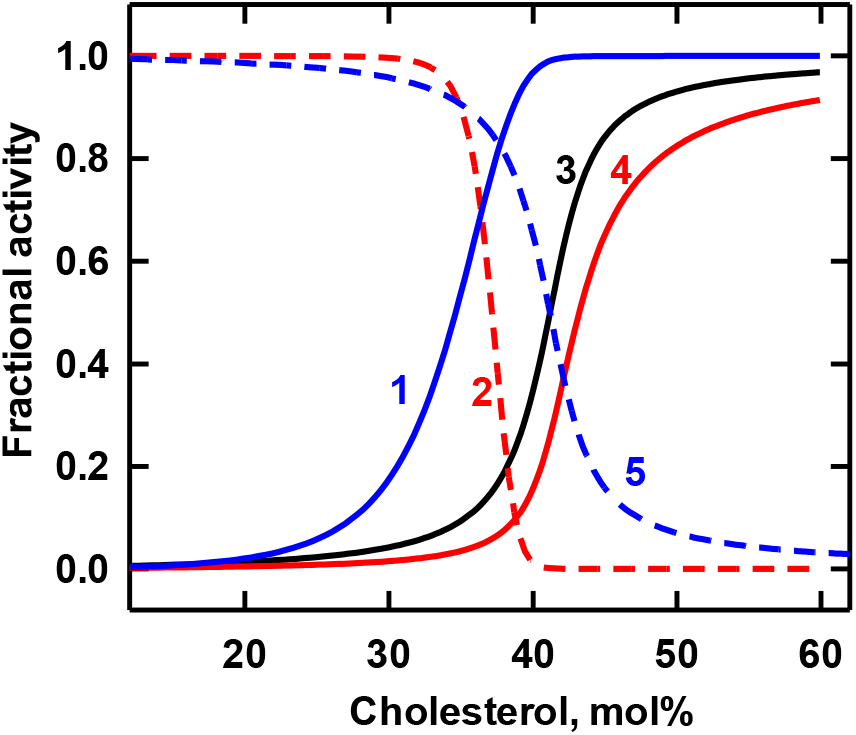
Simulation of the association of cholesterol with five transporters in a hypothetical plasma membrane. Using the values obtained above for K_Ln_ and n, we simulated sterol binding curves for the proteins in their native environment, the plasma membrane. The bilayer was represented by a 1:1 mixture of two phospholipids with stoichiometries of C:P = 1:1 and 1:2 and sterol association constants of 5,000 and 2,500, respectively.^24, 25^ The stoichiometric equivalence point of such bilayers is 43 mol% cholesterol. The values for the affinities of the proteins are given in Table 7.

**Table 7.** Values used in Figure 7.

| Curve | Protein | $n$ | $K_{Ln}$ | Code |
| --- | --- | --- | --- | --- |
| 1 | AChR | 2 | $9 \times 10^5$ | Blue line |
| 2 | BK | 4 | $1 \times 10^{11}$ | Red dash |
| 3 | GAT | 1 | 100 | Black line |
| 4 | Kir 3.4* | 1 | 35 | Red line |
| 5 | Kir2 | 1 | 100 | Blue dash |

Endomembranes (i.e., the cytoplasmic organelles taken as a whole) appear to have a very low cholesterol content (namely, ≤ 10 mol%) and a very low cholesterol affinity.^24, 25^ It was of interest to simulate the association of the integral membrane proteins with cholesterol in such an environment. Figure 8 suggests that proteins with modest cholesterol affinity (namely, those characterized in Figures 1-3) would be partially filled with cholesterol in the weakly-avid intracellular compartment while the oligomers with high affinity and cooperativity would be essentially saturated. Of course, the cholesterol content of various organelles such as the endoplasmic reticulum and distal Golgi membranes differ, and the maturation and modifications of these proteins presumably varies among them. Thus, this exercise is simply heuristic.

**Figure 8.**
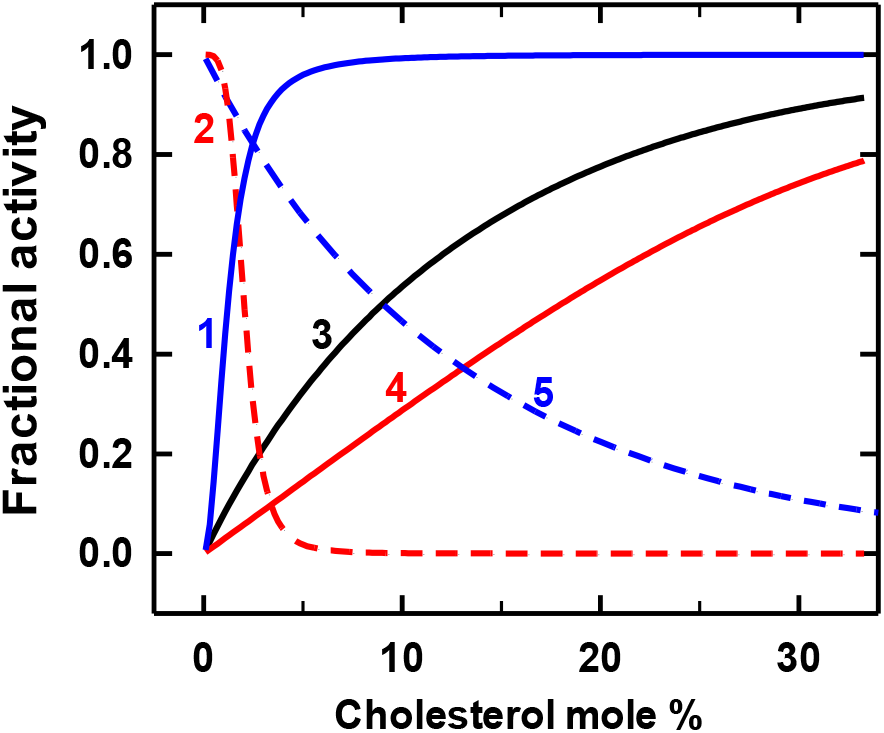
Simulation of the association of cholesterol with proteins in membranes with the phospholipid characteristics of hypothetical endomembranes. We used the values for K_Ln_ and n for five of the transporters studied above and endomembranes that contained a 30/70 mixture of two phospholipids with stoichiometries of C:P = 1:1 and 1:2 and sterol association constants K_P_ = 21 and 10, respectively.^24^ The stoichiometric equivalence point of such bilayers is 39 mol% cholesterol The characteristics of the proteins are given in Table 8.

**Table 8.** Values used in Figure 8.

| Curve | Protein | $n$ | $K_{Ln}$ | Code |
| --- | --- | --- | --- | --- |
| 1 | AChR | 2 | $9 \times 10^5$ | Blue line |
| 2 | BK | 4 | $1 \times 10^{11}$ | Red dash |
| 3 | GAT | 1 | 100 | Black line |
| 4 | Kir3.4* | 1 | 35 | Red line |
| 5 | Kir2 | 1 | 100 | Blue dash |

## DISCUSSION AND CONCLUSIONS

Cholesterol affects the structure and function of a variety of plasma membrane proteins (see Introduction). As illustrated by the figures, the activity of these proteins can have a deeply sigmoidal dependence on membrane cholesterol concentration. These thresholds will reflect both the cooperative binding of the sterol by oligomeric proteins and competition by membrane phospholipids for cholesterol.^46^ Here, we have explored a computational model that estimates the cholesterol association constants and binding stoichiometries of integral membrane proteins based on their competition for the sterol with the membrane phospholipids.

The approach was tested by matching simulations to the cholesterol dependence curves for the activities of six integral membrane proteins. GAT, Kir2 and Kir3.4* did not appear to bind cholesterol cooperatively; that is, n = 1 gave the best matches in Figures 1-3. Even so, these proteins could have bound multiple cholesterol molecules non-cooperatively. The deep sigmoidicity of the isotherms suggests that their sterol affinities are weak (namely, K_L1_ = 35-100) compared to those of typical phospholipids, the K_P_ values of which are in the range of 100-5,000.^29^ Figure 7 suggests that the cholesterol dependence of these proteins would have a threshold near stoichiometric equivalence with the plasma membrane phospholipids, ∼43 mol%. This is the homeostatically-regulated physiologic setpoint of plasma membrane cholesterol.^24^ Thus, as previously proposed, such proteins might have evolved to respond acutely to small variations in the concentration of plasma membrane cholesterol at its resting level.^30^

The data for the other three oligomers were not well fit using n = 1 (Figures 4-6). Rather, they appeared to bind the sterol cooperatively. Furthermore, their subunit association constants were roughly an order of magnitude greater than those of the three non-cooperative proteins (Table 9). Figure 7 suggests that this form of the BK channel and the acetylcholine receptor are saturated with cholesterol at the physiological setpoint of the plasma membrane; all else being equal, they might therefore be insensitive to small variations in its sterol concentration in vivo.

**Table 9.**
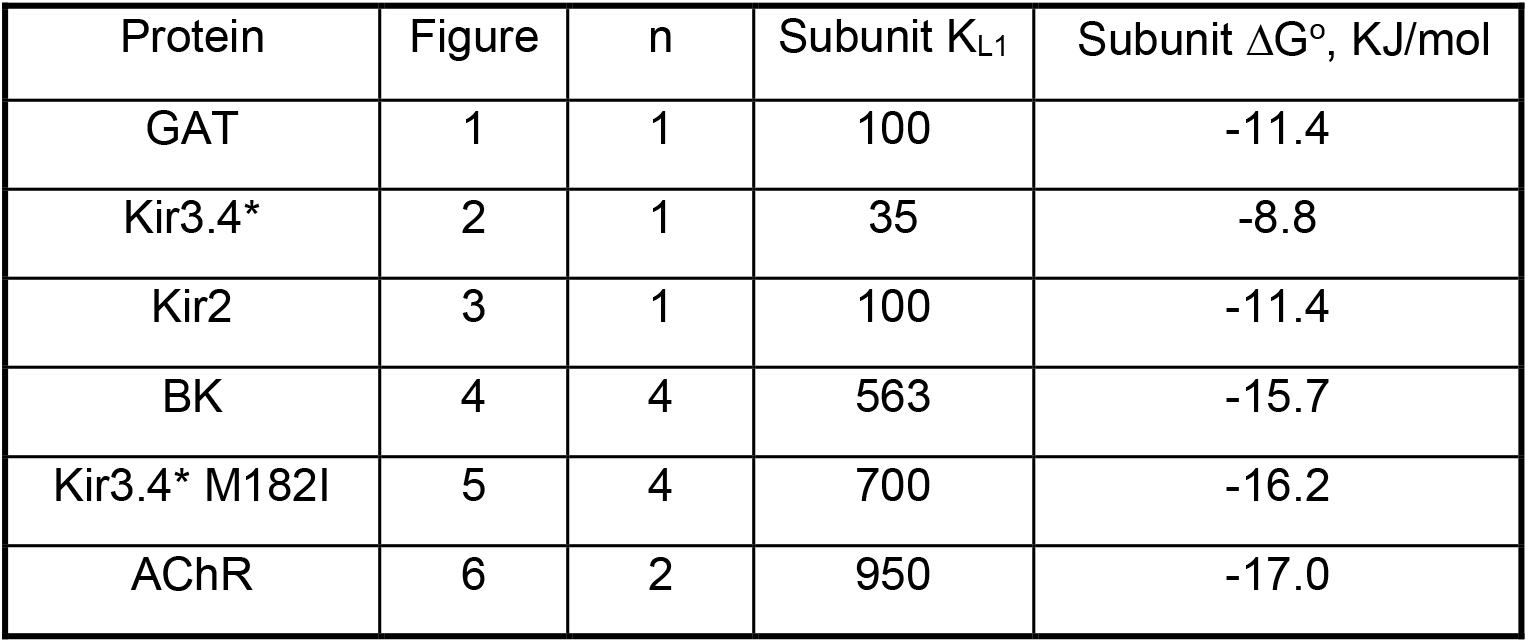
Cholesterol binding characteristics estimated for six transporters.

| Protein | Figure | n | Subunit $K_{L1}$ | Subunit $\Delta G^\circ$, KJ/mol |
| --- | --- | --- | --- | --- |
| GAT | 1 | 1 | 100 | -11.4 |
| Kir3.4* | 2 | 1 | 35 | -8.8 |
| Kir2 | 3 | 1 | 100 | -11.4 |
| BK | 4 | 4 | 563 | -15.7 |
| Kir3.4* M182I | 5 | 4 | 700 | -16.2 |
| AChR | 6 | 2 | 950 | -17.0 |

Table 9 gives the free energy changes inferred for the association of cholesterol with the six transporters. The values are in the middle of a range of estimates (from less than -5 to greater than -50 KJ/mol) computed by different methods for multiple binding sites on a variety of integral membrane proteins.^14–21, 45^ Notably, the cluster of ΔG° values obtained here (namely, - 8.8 to -17.0 KJ/mol) fall in the range of those determined for G protein coupled receptors by molecular dynamics and docking methods (namely, ΔG° = -12.9 to -20.0 KJ/mol); see the Supplement to Lee.^20^

The present analysis has limitations. (a) The experimental data sets were sparse, undermining the precision of the estimates of the sterol association constants and, especially, the binding stoichiometries which depend strongly on the contour of the isotherm. In this model, however, the curves computed for all binding stoichiometries of a protein intersect at the same membrane cholesterol concentration and therefore have the same K_L1_ value (see Methods). Consequently, good estimates of the sterol affinity of the subunit of an oligomer can be secured even when its stoichiometry is uncertain. (b) The model may be too simple, given the complex behavior of many membrane proteins.^47^ For example, a protein might not be an ideal oligomer with n values that are integers, as assumed here. Also, the model stipulates that the only effect of the bilayer on the binding of the sterol to the protein is through competition by the phospholipids while, in reality, other kinds of interactions may obtain.^4^ On the other hand, the matches of simulation to data were satisfactory without the need for a more elaborate model. (c) In vitro may not replicate in vivo. For example, the nearly complete inhibition by cholesterol of the type of BK channel studied in Figure 4 might not obtain in vivo where factors such as cytoplasmic calcium, membrane voltage and accessory proteins serve as physiologic regulators.^38, 40^ In this regard, the model could be useful in comparing values obtained for a plasma membrane protein *in vitro* and *in vivo*. (d) The analysis does not enumerate individual sterol binding sites on a protein or report on dynamics or other molecular features. Rather, it characterizes the equilibrium binding reaction that affects protein activity. (e) The K_Ln_ values obtained depend on characteristics assumed for the phospholipid mixtures. Only approximations are available for some of these.^29^ This might not be a big problem, however. For example, doubling or halving the K_P_ value assigned to curve 3 in Figure 1 (namely, changing the K_P_ from 200 to 100 or 400) yielded K_L1_ values of 55 and 190 rather than the best match, K_L1_ = 100. The flanking K_L1_ estimates correspond to ρG° values of -9.9 and -13.0, not far from the best match: ρG° = -11.4 KJ/mol. Thus, the K_Ln_ values we have obtained should be meaningful despite their uncertain accuracy. (f) Could the experimental sigmoidal isotherms simply reflect cooperative cholesterol binding by the protein without competition by the phospholipids? Some of our simulations were consistent with this supposition (not shown). However, the extensive evidence for sterol-phospholipid complexation makes the central premise of the model secure.^22,23^

The model should be applicable to any integral membrane protein or other ligand that binds cholesterol; for example, Patched and Scap-Insig.^20, 48^ Future applications will surely deliver better estimates of K_Ln_ and n by analyzing more detailed sterol-dependence isotherms. Liposomes, planar bilayers, intact cells and other membrane preparations can be studied. It would be best to employ synthetic bilayers containing a single phospholipid species with a stoichiometry of C:P = 1:1 and a well-determined cholesterol affinity; for example, 1-palmitoyl-2-oleoyl-*sn*-glycero-3-phosphocholine (POPC).^29^ For the study of plasma membrane proteins in situ, the characteristics of erythrocyte phospholipids can be used as the reference, as in Figure 3.^24, 29^ The model posits that K_Ln_ and n will not vary from one bilayer environment to another; this is a testable premise. For example, the influence of charged phospholipids can be examined. In this regard, the values obtained for proteins in their native plasma membranes might reflect the influence of physiological factors—so, how they differ from those obtained using synthetic bilayers should be informative. The method not only enables the study of individual proteins in different environments but also the comparison of a variety of proteins under identical conditions. Such information might shed light on why these proteins have evolved to associate with sterols in the first place.

## ASSOCIATED CONTENT

### Supporting Information

The Supporting Information is available free of charge. It contains a duplicate description of the model, the computational code and the derivation of Eqs.(5) and (6).

## ACKNOWLEDGMENTS

The authors thank Francisco Barrantes (Pontifical Catholic University of Argentina), Anna Bukiya (University of Tennessee Health Science Center), Irena Levitan (University of Illinois at Chicago) and Eduardo Perozo (University of Chicago).

## Notes

The authors declare no competing financial interest.

## Notes

### Competing Interest Statement

The authors have declared no competing interest.

### Summary of Updates

Clarification of presentation.

